# Replication and characterization of *CADM2* and *MSRA* genes on human behavior

**DOI:** 10.1101/110395

**Authors:** Brian Boutwell, David Hinds, the 23andMe Research Team, Jorim Tielbeek, Ken K. Ong, Felix R. Day, John R.B. Perry

## Abstract

Progress identifying the genetic determinants of personality has historically been slow, with candidate gene studies and small-scale genome-wide association studies yielding few reproducible results. In the UK Biobank study, genetic variants in *CADM2* and *MSRA* were recently shown to influence risk taking behavior and irritability respectively, representing some of the first genomic loci to be associated with aspects of personality. We extend this observation by performing a personality “phenome-scan” across 16 traits in up to 140,487 participants from 23andMe for these two genes. Heritability estimates for these traits ranged from 5-19%, with both *CADM2* and *MSRA* demonstrating significant effects on multiple personality types. These associations covered all aspects of the big five personality domains, including specific facet traits such as compliance, altruism, anxiety and activity / energy. This study both confirms and extends the original observations, highlighting the role of genetics in aspects of mental health and behavior.

## Main text

Progress identifying the individual genes underpinning the heritable component (h2=40-60%)(1) of psychiatric disease and personality has lagged behind that of other complex traits(2). A number of reasons might explain this lag, including strong evolutionary selection against common alleles with large effects, a lack of large studies with detailed and harmonized trait information, disease misclassification and subjective clinical criteria. Candidate gene studies have yielded few, if any, credible associations that are replicated(2)(3). In the last few years, suitably powered genome-wide association study (GWAS) meta-analyses have begun identifying handfuls of genetic variants associated with severe psychiatric disease(4)(5). This success, alongside the arrival of large studies such as UK Biobank(6), prompted the hunt for alleles associated with other behavioral and psychological outcomes.

Notably, a recent GWAS on educational attainment(7) identified 74 robustly associated genomic loci, which were subsequently demonstrated to have a causal effect on longevity(8). We previously reported some of the first genomic loci to be robustly associated with aspects of personality – variation in *CADM2* with risk taking propensity and *MSRA* with irritability(9). These variants had a demonstrable effect on reproductive choice and fecundity, yet the breadth of behavioral traits they were associated with remained unclear. This current analysis replicates and extends these prior associations by examining 16 personality constructs using data from 23andMe, Inc., a personal genetics company.

## Methods

### Human Subjects

All 23andMe research participants included in the analysis provided informed consent and answered surveys online according to a human subjects protocol approved by Ethical & Independent Review Services, a private institutional review board.

### Personality Inventories

Soto and John(10) proposed 10 facet scales based on global Big Five Inventory (BFI) items (i.e., two facet scales for each domain captured in the BFI). We computed the 10 facet scales, as well as the traditional BFI categories, using data collected from a web-based version of the Big Five Inventory. Separately, we used a single question to assess risk comfort (see supplementary materials for additional information). Items were coded such that higher scores on each dimension corresponded to the increased presence of a particular trait (i.e., higher scores on the “compliance” items represent more compliant and agreeable tendencies). All phenotypes were coded on a 5-point scale according to the answers given to the personality questions listed in Supplementary data. For phenotypes based on more than one question an individual’s score is the average of their answers across questions.

### Genetic analysis

Genotyping of research participants was performed as previously described(11). Genotypes for rs1865251 and rs658385 were imputed (minimum imputation quality > 0.95) using the March 2012 “v3” release of 1000 Genomes reference haplotype panel. Genetic association results were obtained from linear regression models assuming additive allelic effects. These models included as covariates – age, sex and the top five genetically determined principal components to account for population structure. The reported SNP association test P-values were computed from likelihood ratio tests.

SNP-based heritability was estimated using LD score regression, implemented in the LDSC software package. Default software parameters were used and only common SNPs (MAF>5%) present in HapMap 3 were considered.

## Results

Phenotypes and genotypes were available in up to 140,487 research participants from 23andMe, independent from the previous study. A total of 16 personality constructs were defined on the basis of answers from 45 questions (**Supplementary Table 1**). We estimated a significant SNP-based heritability for each of these 16 traits (**Figure 1**), ranging from 5.1% (risk taking) to 19.1% (extraversion).

**Figure 1.**
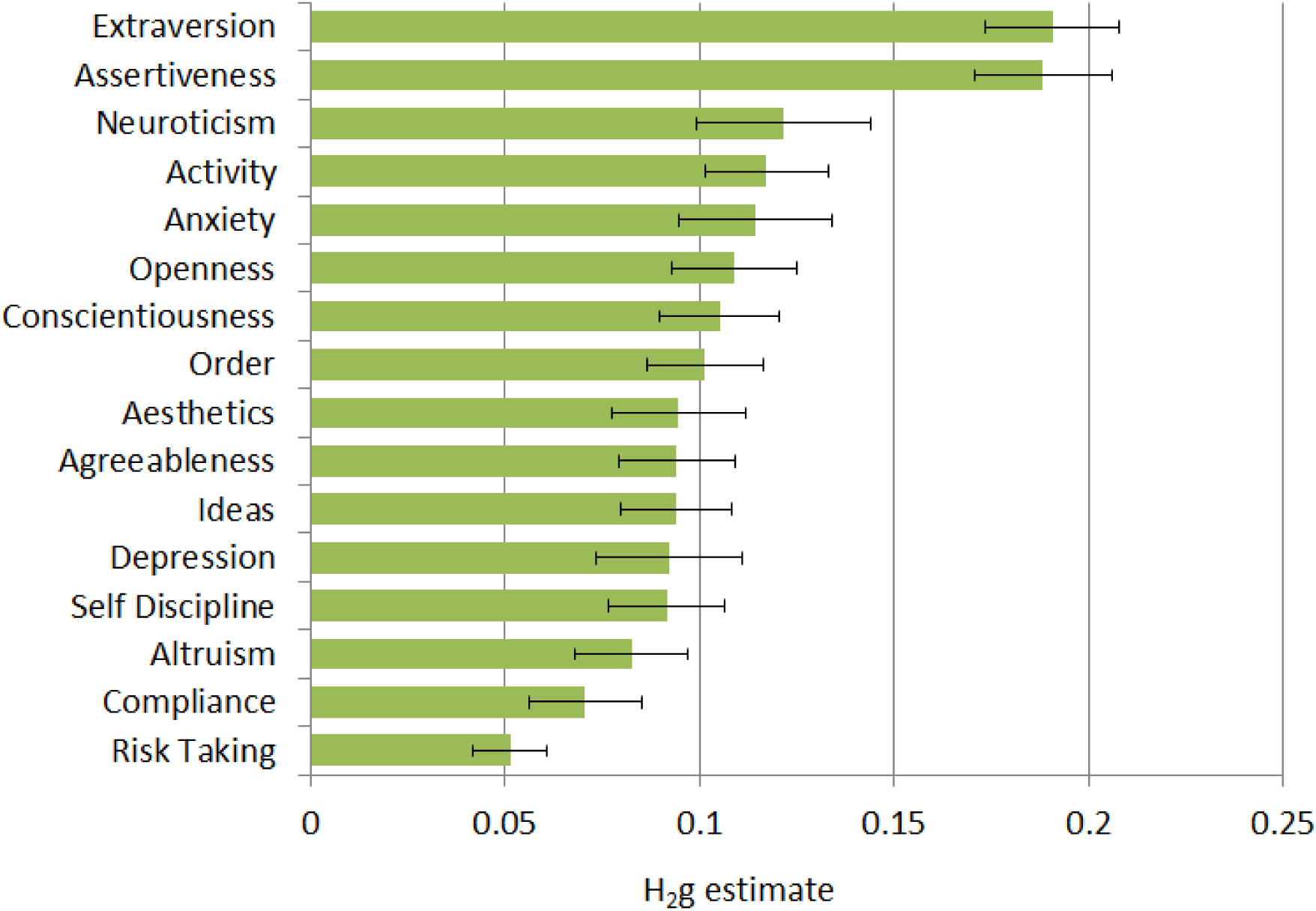
Genome-wide SNP-based heritability estimates for the 16 personality traits tested.

The *CADM2* risk taking allele (best Hapmap2 proxy = rs1865251, “C” allele) was significantly associated (P=1.6×10^-5^) with outcome to the question “Overall, do you feel comfortable or uncomfortable taking risks?”, confirming the previous observation. An additional nine personality traits exhibited a nominal (P<0.05) association (**Figure 2**), the most significant of which was reduced anxiety levels (P=8.6×10^-6^).

**Figure 2.**
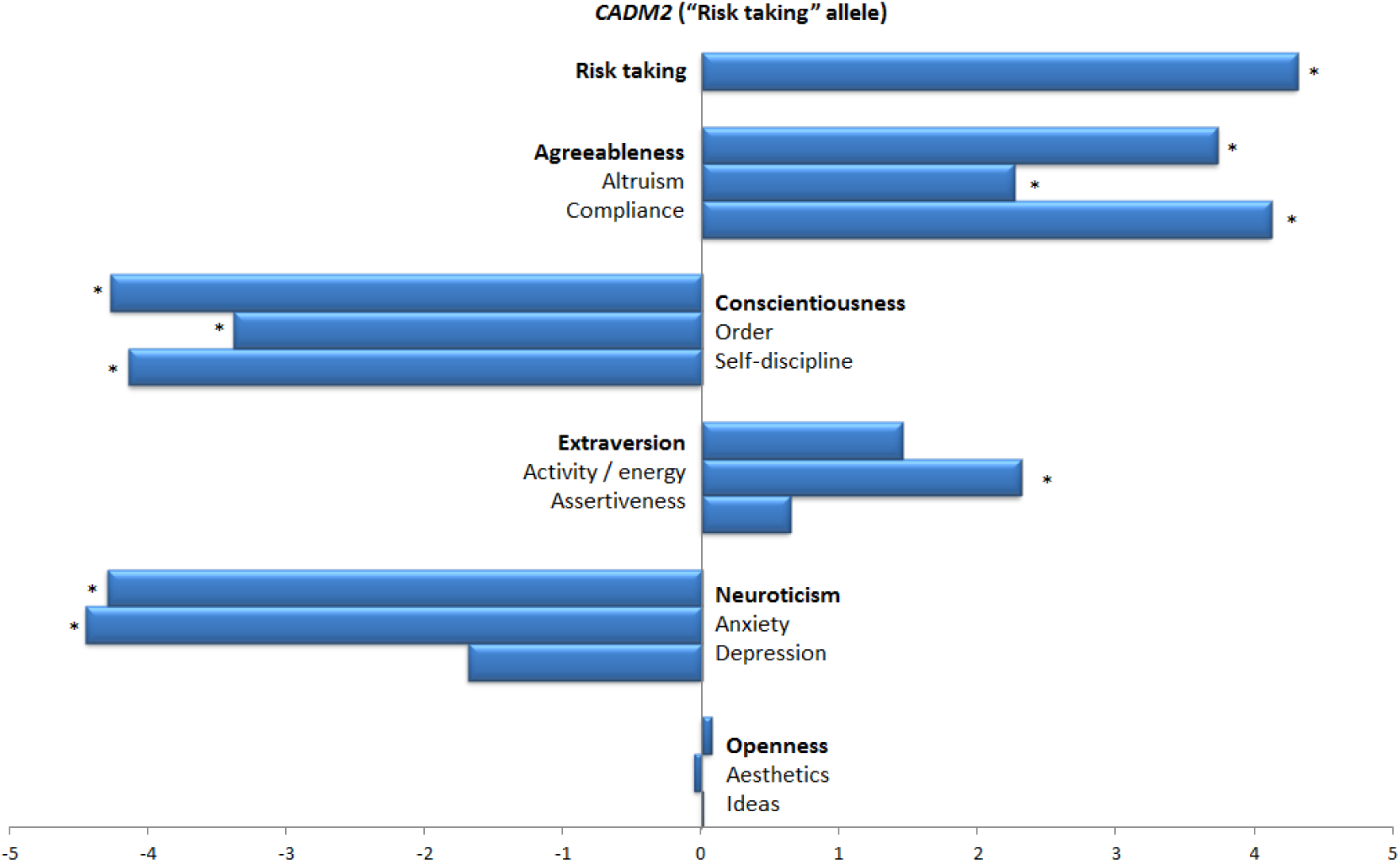
Personality types associated with the *CADM2* risk taking allele. Plot shows signed Z score association statistic for the “C” allele of rs1865251. Personality traits that are nominally significant (Z= +/- 1.96 ~ P<0.05) are highlighted with an asterisk (*). The 10 facet traits are organized within their corresponding “big five” personality domains (shown in bold).

For *MSRA*, there was no single personality trait that matched self-reported irritability in the 23andMe dataset; however, the allele previously associated with increased irritability was significantly associated (**Figure 3**) with reduced energy and enthusiasm (“activity”, P=7.2×10^-4^), less compliance (P=4.8×10^-4^), more depressive feelings (P=6×10^-7^), more neuroticism (P=1.1×10^-7^), less creative thinking (“ideas”, P=0.02), more introversion (P=8×10^-3^), more anxiousness (P=6.9×10^-5^), less openness to new ideas (P=9.8×10^-3^) and more adverse feelings to taking risks (P=0.01).

**Figure 3.**
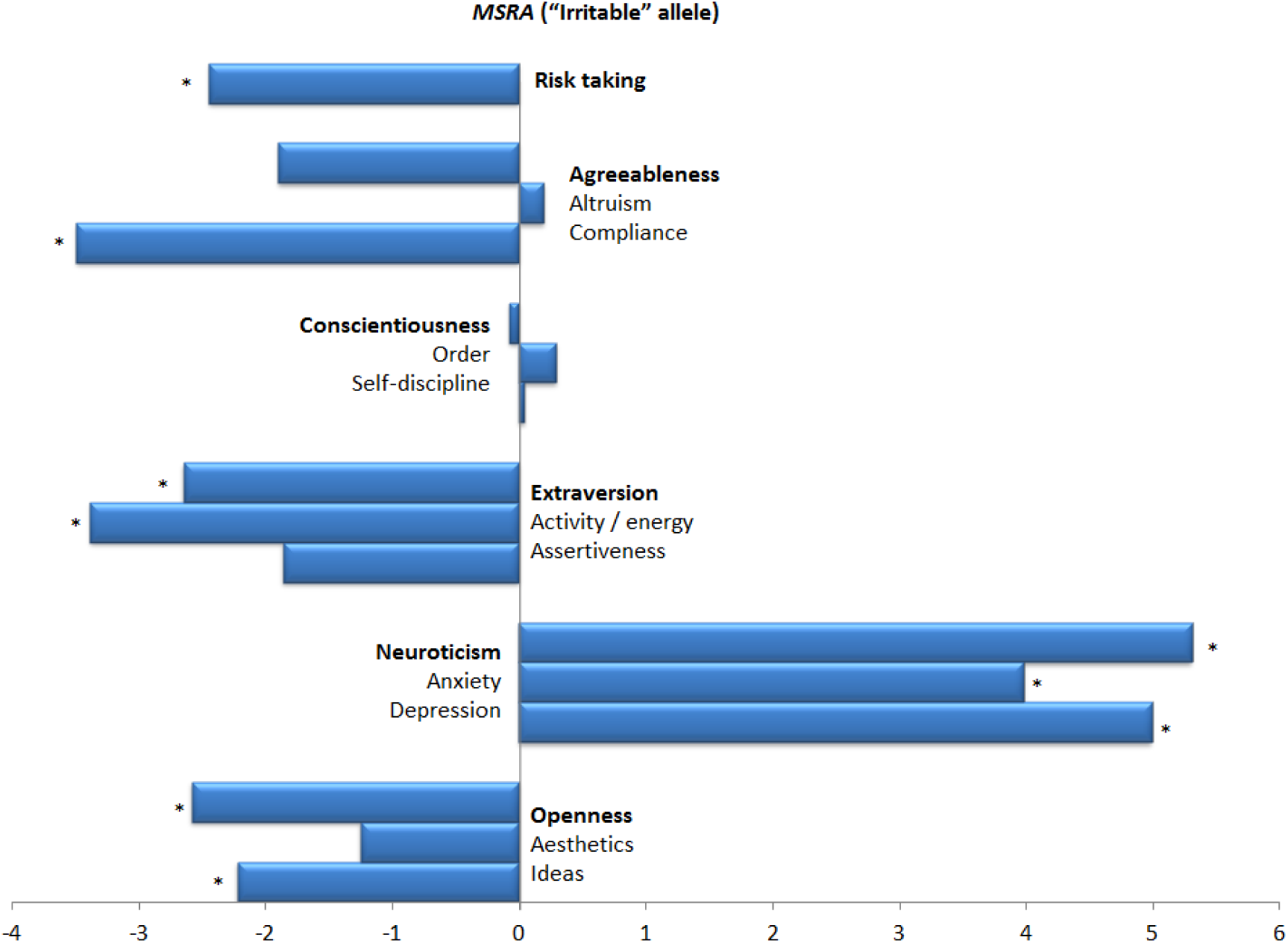
Personality types associated with the *MSRA* increased irritability allele. Plot shows signed Z score association statistic for the “T” allele of rs658385. Personality traits that are nominally significant (Z= +/- 1.96 ~ P<0.05) are highlighted with an asterisk (*). The 10 facet traits are organized within their corresponding “big five” personality domains (shown in bold).

## Discussion

Large-scale GWAS meta-analyses are now routinely comprised of hundreds of thousands of samples, and as a result possess the requisite statistical power for elucidating the genetic basis of complex human personality traits. Within the last year, several studies have identified the first robustly associated genetic variants associated with these outcomes, with a handful of loci discovered for subjective well-being(12), neuroticism(12, 13), extraversion(13) and conscientiousness(13). In addition, we previously reported genetic variation in *MSRA* and *CADM2* linked to a range of behavioral reproductive outcomes(9). Here, the signal in *CADM2* was initially linked to a single self-reported behavioral outcome – risk taking propensity – in the UK Biobank study. The risk taking allele was associated with earlier age at first sex, increased number of sexual partners, a higher number of offspring, increased body mass index and decreased cognitive processing speed. Our current study replicates and extends this observation, demonstrating that this risk taking behavior increasing allele is broadly associated with a “care free” and optimistic personality type. *CADM2* is widely expressed in a number of neuronal tissues and implicated in glutamate signaling, γ-aminobutyric acid transport and neuron cell–cell adhesion(14). In contrast, the *MSRA* allele promoting later age at first sex was first demonstrated to have an association with increased self-reported irritability. We extend this observation to portray a personality type dominated by anxiety, neuroticism, depression, lack of compliance and reduced energy levels. Further experimental work will be required to identify the likely effector gene behind this association, given the extensive linkage disequilibrium in the region. Since our initial observation this locus has been reported at genome-wide significance for neuroticism(12, 13), however our current result demonstrates a broader personality influence.

In summary, our study and others demonstrates that genetic studies are now suitably powered to identify reproducibly associated genetic determinants of personality. Genetic correlation analyses have previously demonstrated significant overlap in the genes responsible for “normal” personality variation and those involved in severe psychiatric disease(13). This suggests that expanded personality studies may eventually provide complementary insight into the etiology of severe disease states, when a more substantial fraction of the heritability of personality type is established. It also paves the way for the understanding of causes and consequences of these conditions through the use of Mendelian randomization and similar methods.

## Acknowledgements

We thank the customers of 23andMe who consented to participate in research and the employees of 23andMe who together have helped made this research possible.

### Author List for the 23andMe Research Team

Michelle Agee, Babak Alipanahi, Adam Auton, Robert K. Bell, Katarzyna Bryc, Sarah L. Elson, Pierre Fontanillas, Nicholas A. Furlotte, David A. Hinds, Bethann S. Hromatka, Karen E. Huber, Aaron Kleinman, Nadia K. Litterman, Matthew H. McIntyre, Joanna L. Mountain, Carrie A.M. Northover, J. Fah Sathirapongsasuti, Olga V. Sazonova, Janie F. Shelton, Suyash Shringarpure, Chao Tian, Joyce Y. Tung, Vladimir Vacic, Catherine H. Wilson

## References

1. Polderman, T.J.C., Benyamin, B., de Leeuw, C.A., Sullivan, P.F., van Bochoven, A., Visscher, P.M. and Posthuma, D. (2015) Meta-analysis of the heritability of human traits based on fifty years of twin studies. Nat. Genet., 47, 702–709.

2. Chabris, C.F., Lee, J.J., Cesarini, D., Benjamin, D.J. and Laibson, D.I. (2015) The Fourth Law of Behavior Genetics. Curr. Dir. Psychol. Sci., 24, 304–312.

3. Munafo, M.R., Clark, T.G., Moore, L.R., Payne, E., Walton, R. and Flint, J. (2003) Genetic polymorphisms and personality in healthy adults: A systematic review and meta-analysis. Mol. Psychiatry, 8, 471–484.

4. Ripke, S., Neale, B.M., Corvin, A., Walters, J.T.R., Farh, K.-H., Holmans, P.A., Lee, P., Bulik-Sullivan,B.,Collier, D.A. and Huang, H. (2014) Biological insights from 108 schizophrenia-associated genetic loci. Nature, 511, 421.

5. Lee, S.H., Ripke, S., Neale, B.M., Faraone, S. V., Purcell, S.M., Perlis, R.H., Mowry, B.J., Thapar, A., Goddard, M.E., Witte, J.S., et al. (2013) Genetic relationship between five psychiatric disorders estimated from genome-wide SNPs. Nat. Genet., 45, 984–94.

6. Sudlow, C., Gallacher, J., Allen, N., Beral, V., Burton, P., Danesh, J., Downey, P., Elliott, P., Green, J., Landray, M., et al. (2015) UK Biobank: An Open Access Resource for Identifying the Causes of a Wide Range of Complex Diseases of Middle and Old Age. PLoS Med., 12, e1001779.

7. Okbay, A., Beauchamp, J.P., Fontana, M.A., Lee, J.J., Pers, T.H., Rietveld, C.A., Turley, P., Chen, G.-B., Emilsson, V., Meddens, S.F.W., et al. (2016) Genome-wide association study identifies 74 loci associated with educational attainment. Nature, 533, 539–542.

8. Marioni, R.E., Ritchie, S.J., Joshi, P.K., Hagenaars, S.P., Okbay, A., Fischer, K., Adams, M., Hill, D.W., Davies, G., Social Science Genetic Association Consortium, et al. (2016) Genetic variants linked to education predict longevity. 10.1073/pnas.1605334113.

9. Day, F.R., Helgason, H., Chasman, D.I., Rose, L.M., Loh, P.-R., Scott, R.A., Helgason, A., Kong, A., Masson, G., Magnusson, O.T., et al. (2016) Physical and neurobehavioral determinants of reproductive onset and success. Nat. Genet., advance on, 617–623.

10. Soto, C.J. and John, O.P. (2009) Ten facet scales for the Big Five Inventory: Convergence with NEO PI-R facets, self-peer agreement, and discriminant validity. J. Res. Pers., 43, 84–90.

11. Day, F.R., Hinds, D.A., Tung, J.Y., Stolk, L., Styrkarsdottir, U., Saxena, R., Bjonnes, A., Broer, L., Dunger, D.B., Halldorsson, B. V, et al. (2015) Causal mechanisms and balancing selection inferred from genetic associations with polycystic ovary syndrome. Nat. Commun., 6, 8464.

12. Okbay, A., Baselmans, B.M.L., De Neve, J.-E., Turley, P., Nivard, M.G., Fontana, M.A., Meddens, S.F.W., Linnér, R.K., Rietveld, C.A., Derringer, J., et al. (2016) Genetic variants associated with subjective well-being, depressive symptoms, and neuroticism identified through genome-wide analyses. Nat. Genet., 10.1038/ng.3552.

13. Lo, M.-T., Hinds, D.A., Tung, J.Y., Franz, C., Fan, C.-C., Wang, Y., Smeland, O.B., Schork, A., Holland, D., Kauppi, K., et al. (2016) Genome-wide analyses for personality traits identify six genomic loci and show correlations with psychiatric disorders. Nat. Genet., 10.1038/ng.3736.

14. Ibrahim-Verbaas, C. a, Bressler, J., Debette, S., Schuur, M., Smith, a V, Bis, J.C., Davies, G., Trompet, S., Smith, J. a, Wolf, C., et al. (2015) GWAS for executive function and processing speed suggests involvement of the CADM2 gene. Mol. Psychiatry, 10.1038/mp.2015.37.

